# Loss of essential outer membrane functions causes drug hypersensitization in *Acinetobacter baumannii* overexpressing multidrug efflux pumps

**DOI:** 10.1101/2024.01.04.574119

**Authors:** Efrat Hamami, Wenwen Huo, Katherine Neal, Isabelle Neisewander, Shivangi Fnu, Joel S. Freundlich, Edward Geisinger, Ralph R. Isberg

## Abstract

Elevated expression of Resistance-Nodulation-Cell Division (RND) drug transporters is commonly observed in clinical isolates of *Acinetobacter baumannii*, a nosocomial pathogen associated with multidrug-resistant (MDR) infections. We describe here a CRISPRi platform directed toward identifying essential gene hypomorphs that preferentially change resistance to the fluoroquinolone antibiotic ciprofloxacin in RND pump overproducers. An sgRNA library including single and double nucleotide mutations directed against essential genes of *A. baumannii* was constructed and introduced into multiple strain backgrounds, allowing strain-specific, titratable knockdown efficiencies to be analyzed. Other than NusG depletions, there were few candidates in the absence of drug treatment that showed lowered fitness specifically in strains overexpressing the RND efflux pumps AdeAB, AdeIJK, or AdeFGH. In the presence of ciprofloxacin, the hypomorphs that caused hypersensitivity were predicted to result in outer membrane dysfunction, with the AdeFGH overproducer appearing particularly sensitive to such disruptions. Most notably, depletion of the predicted monovalent cation-proton antiporter component PhaF compromised efflux pump function, as depletion of the antiporter resulted in increased ciprofloxacin accumulation in strains overproducing AdeFGH and disrupted cytosolic pH. On the other hand, depletions of translation-associated proteins and components of the proton-pumping ATP synthase conferred fitness benefits in the presence of the drug in at least two pump-overproducing strains. Therefore, pump overproduction exacerbated stress caused by defective outer membrane integrity, while the activity of at least one efflux pump overproducer required the function of an antiporter that maintains cytosolic pH homeostasis.

**Importance:** *Acinetobacter baumannii* clinical isolates are increasingly multidrug-resistant, leaving patients with few effective treatment options. Many of these isolates are fluoroquinolone-resistant due to drug target mutations in the two major type II topoisomerases, as well as mutations that activate RND efflux pump expression. This work identifies essential gene products that support the fitness of efflux pump hyperexpressers during treatment with the fluoroquinolone antibiotic ciprofloxacin, most of which are involved in OM biogenesis. These findings suggest new strategies for combination therapy with currently available fluoroquinolones to sensitize and combat high-level resistant strains.

## Introduction

*Acinetobacter baumannii* is a gram-negative nosocomial pathogen that causes ventilator-associated pneumonia, bacteremia, and urinary tract infections (1–3). The difficulty in combating diseases caused by this often multidrug-resistant (MDR) organism has stimulated the demand for new antimicrobials and improved therapeutic strategies (1, 3). Among the various determinants of resistance to currently available drugs, hyper-expression of drug efflux pumps is commonly associated with MDR across multiple species (4–9). The Resistance-Nodulation-Division (RND) class of efflux pumps provides the most clinically relevant strategy for extruding antibiotics in *A. baumannii,* with the AdeABC, AdeIJK, and AdeFGH efflux pumps frequently associated with MDR (6–9). Therefore, strategies that target isolates overexpressing these pumps or which hypersensitize these strains to currently available antibiotics are of great interest.

For gram-negative organisms, the primary route to fluoroquinolone resistance (FQR) is via acquisition of mutations in the active sites of the major type II topoisomerases in the cell, DNA gyrase and topoisomerase IV (10). In sequenced *A. baumannii* clinical strains, resistance mutations in both *gyrA* and *parC* are found, encoding the A subunits of gyrase and topo IV, respectively (10). Based on MIC testing and sequencing of isolates, FQR clinical strains also commonly have a third mutation in one of the regulatory genes that activate an RND antibiotic efflux pump. Mutations in these genes, encoded by *adeS*, *adeN*, or *adeL* upregulate the AdeAB(C), AdeIJK, or AdeFGH efflux pumps respectively and result in increased FQR, consistent with fluoroquinolones such as ciprofloxacin being substrates of these efflux systems (6, 7, 9–19).

Previous work identified mutations that caused strain-specific vulnerabilities of pump overproduction in nonessential *A. baumannii* genes. These mutations hindered growth of strains overexpressing efflux pumps or synergized with the fluoroquinolone ciprofloxacin (CIP) to hypersensitize the overproducers to the drug (16). Based on that work, lesions in nonessential genes predicted to alter envelope integrity sensitize efflux pump overproducing strains to CIP treatment. Furthermore, the strain with the activating mutation in *adeL* (causing *adeFGH* overexpression) appeared to result in the most significant physiological consequences, with a broader mutation spectrum found to reduce fitness, and a more robust transcriptional shift was apparent relative to the other efflux pump overproducers. These effects may be due to either special features of this pump or extreme overexpression of the *adeFGH* locus relative to the other pump-overexpressing strains (16).

Most effective antimicrobials interfere with proteins essential for growth, making them desirable targets for identifying mutations that preferentially sensitize strains overproducing efflux pumps. To approach this problem, we used CRISPRi to target essential genes with sgRNAs. The library design takes advantage of a previously developed strategy that generates multiple hypomorphic alleles with varying fitness defects in each essential gene, enabling the discovery of depletions that show strain-specific aggravation of drug sensitivity (20–22).

Depletions that cause drug hypersensitivity were found to predominantly target proteins involved in maintaining outer membrane (OM) integrity. Interestingly, interference of translation and the assembly of the proton pumping F_1_F_0_ ATPase provided the FQR strains with a fitness advantage in the presence of drug. Our work is complementary to other recent studies using CRISPRi to identify essential *A. baumannii* pathways that alter sensitivities to a large panel of antibiotics (23, 24).

## Results

### Development of a CRISPRi strategy to uncover hypomorphs with reduced fitness in FQR strains

In our previous study (16), we identified transposon insertion mutations that reduce the fitness of efflux pump overproducers or that specifically hypersensitize these variants to CIP. Except for very rare exceptions (25), insertion mutations could only be analyzed in nonessential genes. This interfered with our ability to identify proteins encoded by the essential gene set that could similarly sensitize pump overproducers. To overcome this limitation, we devised a CRISPRi screening strategy to identify viable depletion variants with drug- or strain-dependent fitness defects (Fig. 1) (20, 21). We applied a pool design similar to that used for eukaryotic cells (20). To identify essential genes, we inspected Tn-seq data from a previously generated dense transposon bank in the ATCC 17978UN strain background (16, 25, 26) for open reading frames (ORFs) having significantly reduced or no insertions (Materials and Methods (27, 28)). After comparison with other published *A. baumannii* Tn-seq studies, a list of 406 essential genes was generated (16, 26, 29, 30).

**Figure 1:**
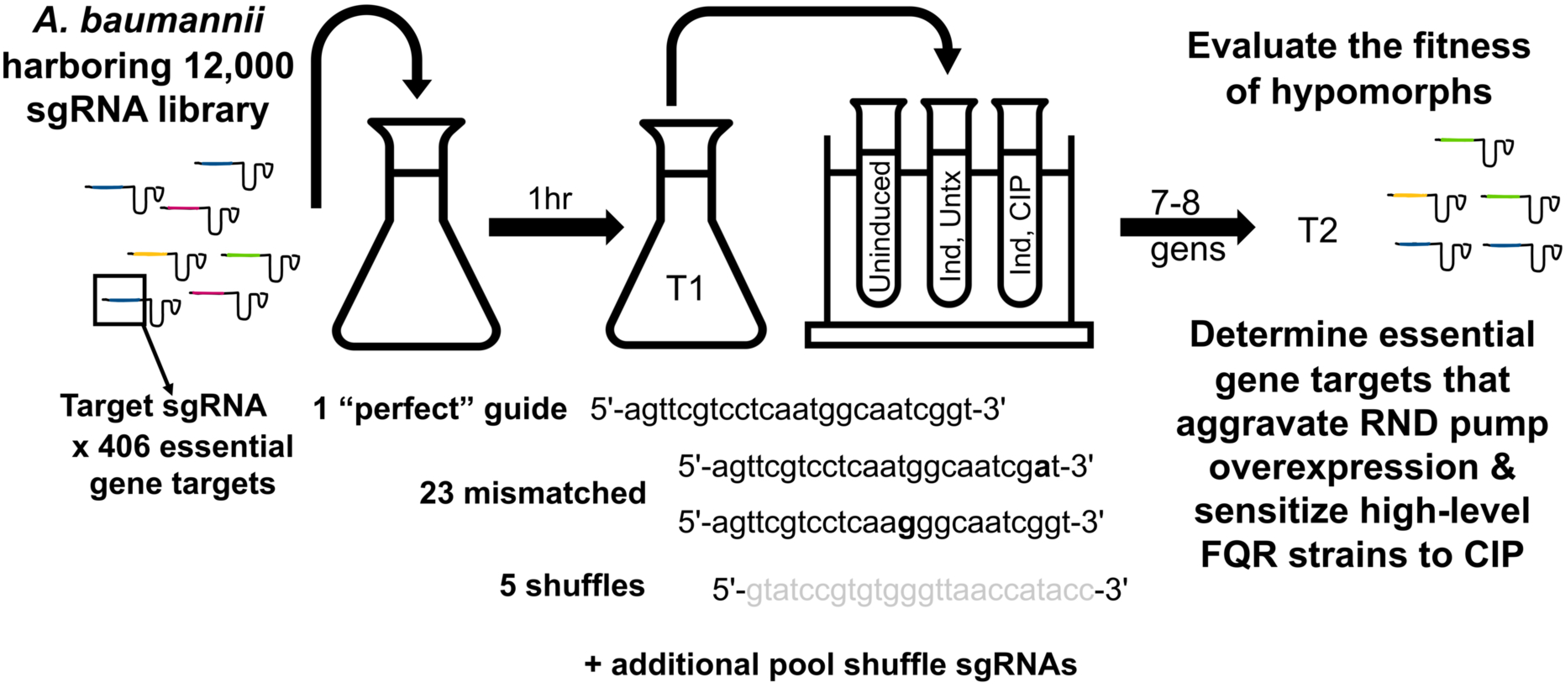
Pooled CRISPRi screen to identify hypomorphs with reduced fitness in FQR strains. Diagram for experimental approach and sgRNA library design (Materials & Methods). sgRNAs were designed to target 406 essential genes in *A. baumannii* (“perfect” guides). For each perfect guide, a set of 23 guides with 1 to 2 mismatched nts were created to achieve titratable knockdown efficiencies as well as 5 sequence-shuffled guides as negative controls. Additional pool shuffle sgRNAs were made to generate a library of 12,000 guides. Cultures were harvested before (T1) and after (T2) outgrowth with or without CIP and the dCas9 inducer aTc. Plasmid was extracted and sequenced to identify hypomorphs with altered fitness under tested conditions. Hypomorph fitness scores, normalized to scrambled guides, were compared between control and experimental groups to determine essential gene targets that generate altered fitness of high-level FQR strains relative to WT in either the presence or absence of CIP. FQR: fluoroquinolone resistant. Ind: dCas9 induced using aTc. Untx: untreated. CIP: ciprofloxacin treated (0.11-0.12 µg/mL for WT, 50 µg/mL for GPN, 55 or 65 µg/mL for GPS, and 60 µg/mL for GPL, Supplementary Data 5).

A single 24-nucleotide (nt) sgRNA was designed for each essential gene by identifying the PAM site on the noncoding strand most proximal to the translational start. For each perfect sgRNA, we devised a set of 23 mismatch sgRNAs that differed from the original guide sequence by 1 or 2 nts at varying locations across the sequence (20). We also created five shuffle sgRNAs comprising the same nts in a disordered fashion to use as negative controls, anticipating that they would have a neutral effect on fitness. Additionally, a collection of approximately 200 shuffle sgRNAs with minimal anticipated genome homology was created, called pool shuffles. The final library consisted of approximately 12,000 sgRNAs with sizes and complexities similar to those of a previously constructed model pool (Fig. 1) (20). To facilitate cloning, the pYDE007 sgRNA delivery vector was converted into a GoldenGate-compatible plasmid to enable efficient construction of the sgRNA library by inserting duplex target oligonucleotides from the pool into the site generated by a Type II restriction enzyme (see Materials and Methods). Control of targeted CRISPRi depletions was provided by an anhydrotetracycline (aTc) inducible dCas9 introduced into the genomes of the WT and FQR pump-overexpressing strains (using a Tn7 strategy originally devised by Peters *et al*. (31) and modified by Bai *et al*. (26); Materials and Methods). The FQR strain panel had the *gyrA parC* (GP) background with either the *adeS*, *adeN,* or *adeL* pump-overproducing mutations (GPS, GPN, and GPL respectively). These strains were then transformed in parallel with the CRISPRi sgRNA plasmid bank, ensuring each pool comprised at least 10^6^ transformants.

### Mismatched sgRNAs result in a gradient of fitness defects

Pools harbored in all strain backgrounds were cultured in LB for 7-8 generations in the absence or presence of aTc or CIP (Materials and Methods; Fig. 1), with cultures harvested before (timepoint T1) and after bank outgrowth (T2). Plasmid DNA was extracted at both timepoints and the sgRNA region amplified from the extracted plasmid DNA, barcoded, and sequenced in parallel (Materials and Methods). Reads were processed to determine the abundance of each unique sgRNA at both time points in all conditions. The sgRNA reads showed a normal distribution (Supp. Fig. 1A) that was maintained after introduction into the WT *A. baumannii* strain (Supp. Fig. 1B). A similar distribution was observed from the shuffle sgRNAs after aTc induction (Supp. Fig. 1C), consistent with nontargeting behavior as expected for negative controls. Fitness was calculated after stoichiometric depletion of target genes across replicate cultures as previously described (Materials and Methods (10, 25, 32), with fitness values normalized to a small set of shuffle sgRNAs that had no statistically significant effect on fitness after dCas9 induction. As anticipated, most shuffle guides showed no change in fitness upon dCas9 induction (Fig. 2A), whereas the majority of sgRNAs with perfect target homology showed diminished fitness upon dCas9 induction (73% of perfect guides reducing fitness based on FDR <0.05, fitness < 0.96 after induction compared to control, with fitness at least 10% lower than uninduced cultures; Fig. 2B). Improved fitness was observed by guides harboring mismatched nts closest to the 3’ PAM site critical for dCas9 targeting (34% of these guides with mismatched nts most proximal to the PAM site reducing fitness, based on FDR <0.05 and fitness < 0.93 after induction, with fitness lowered by at least 10% from uninduced cultures; Fig. 2C) (20, 21). This pattern can be seen by focusing on sgRNA sets targeting individual genes, such as *gyrA*, which encodes subunit A of DNA gyrase, which showed a broad range of fitness defects depending on the mismatch site (Fig. 2D, E).

**Figure 2:**
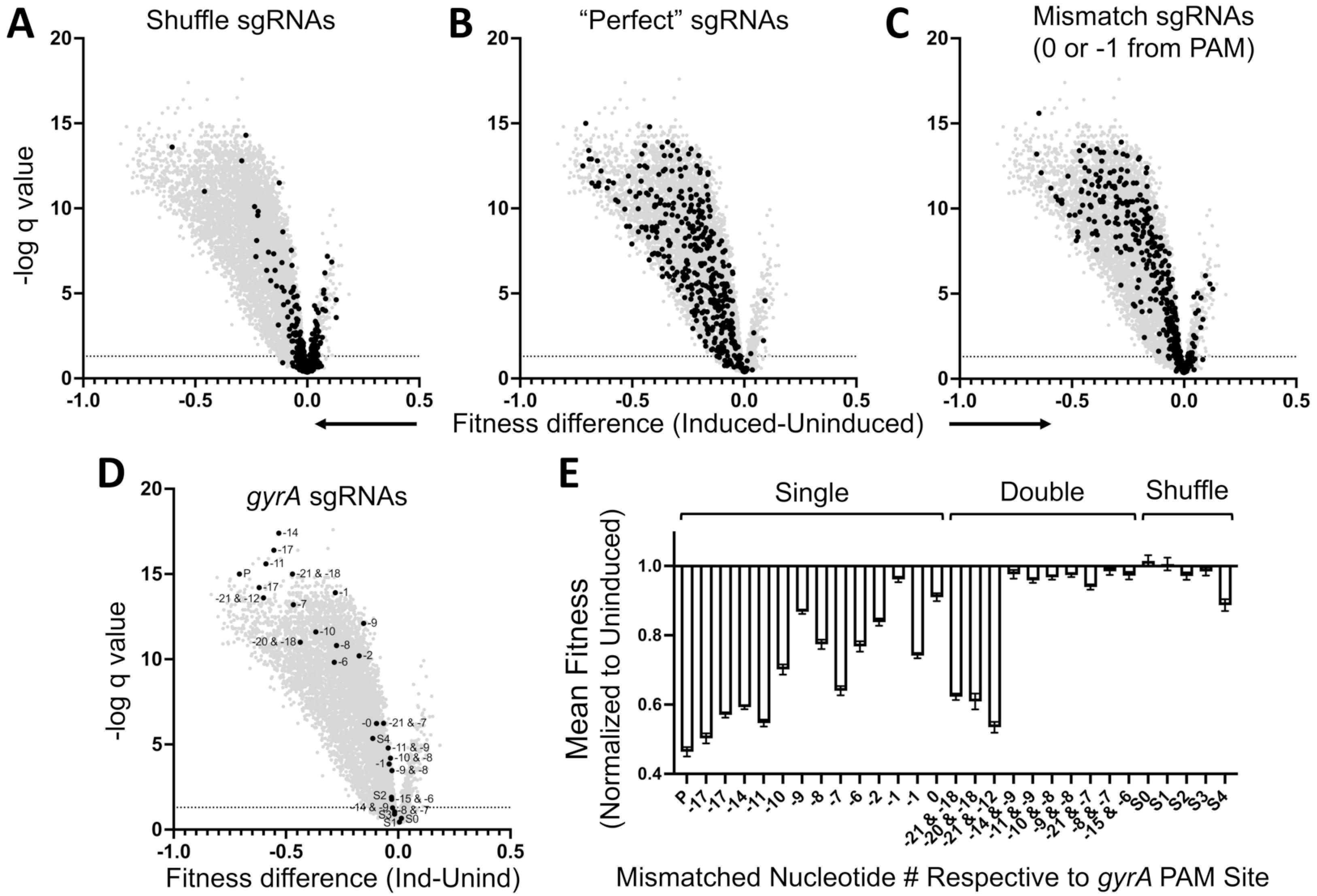
sgRNA fidelity to essential gene sequences and mismatch distance from PAM site correspond with fitness decreases in *A. baumannii* CRISPRi strains. Volcano plots are displayed to show statistical significance as a function of the mean change in fitness scores of hypomorphs in the WT strain background that result from dCas9 induction (100ng/mL aTc, n=10). All symbols refer to unique sgRNAs (light gray). The horizontal dotted line (A-D): FDR or Q=5%. (A) Displayed are shuffle sgRNAs that serve as negative controls (black). (B) The sgRNAs with perfect sequence identity to their target (black). (C) The sgRNAs with mismatched nts located most proximal to the 3’ PAM site (black; nts 23 or 24 of the 24nt sgRNA, -1 or 0 nt from PAM site respectively). (D, E): The sgRNAs designed for *gyrA* (black). S: Shuffle sgRNAs. P: Perfect guide. Positions of mismatched nts in relation to the 3’ PAM site noted, with numbering starting at 0 for the nt immediately upstream the PAM site. (D) Display of mean fitness difference of *gyrA* sgRNAs after induction vs. uninduced. (E) Display of *gyrA* data, showing average fitness scores normalized to fitness in the absence of dCas9 inducer for the perfect sgRNA and sgRNAs with single and double mismatches as well as shuffle sgRNAs (± SEM).

The mild effects observed in the remaining 27% of perfect-match sgRNAs could be due to a variety of factors, including the relative distance of the target sequence from the translation start codon. A distinct relationship was difficult to establish, except when comparing fitness yielded by guides targeting sites 25-40 nts away from the target codon start site versus those recruited to sites only 1-10 nts away (Supp. Fig. 2A-D). As many sgRNAs with high fitness likely resulted from poor target depletion (ex. Supp. Fig. 2E), we determined if continued growth for additional generations (14-16 generations vs. 7-8) could amplify fitness defects. We found that continued growth expanded the number of perfect sgRNAs causing a significant defect (a 35-38% boost in guides with fitness lowered by at least 0.5; Supp. Fig. 3) without a substantial loss of fitness for the shuffle guides (Supp. Fig. 4). These data indicate that the defects observed in hypomorphs can be aggravated by challenging the sgRNA banks for longer periods of time, consistent with the target proteins being subjected to turnover and dilution through bacterial division.

**Figure 3:**
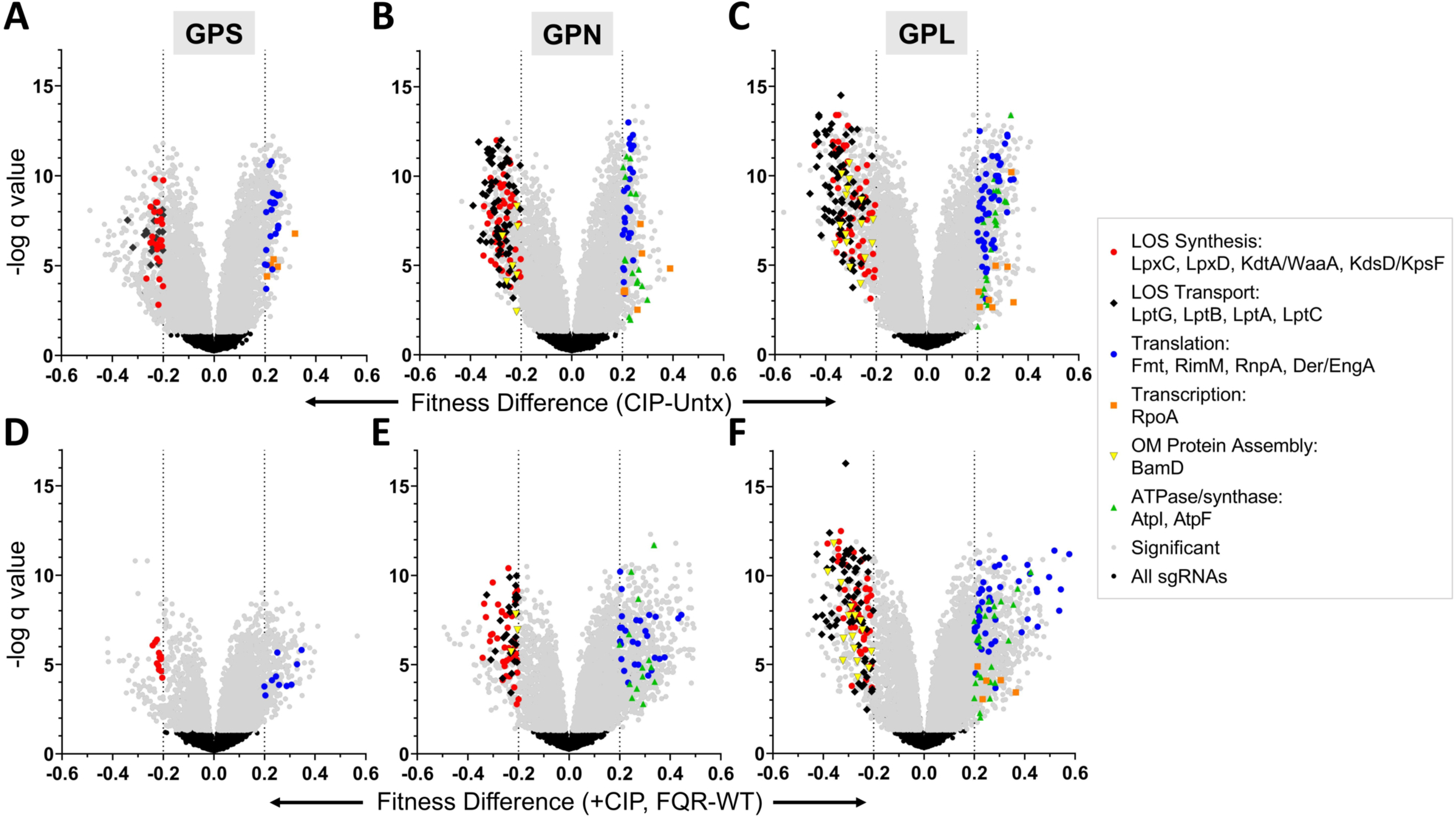
Opposing effects of translation/ATP generation and LOS biogenesis on the fitness of FQR strains relative to WT in the presence of CIP. (A-C) Statistical significance plotted as a function of mean fitness score differences (n=10) between CIP and Untx conditions (top) or FQR and WT strains during CIP treatment (bottom), of the FQR strain backgrounds GPS (A, D), GPN (B, E), or GPL (C, F). Gray: Statistically significant sgRNAs passing the FDR of 5% (33). 16 prominent targets from the CIP CRISPRi screen are highlighted. Dotted lines: selection parameters setting fitness shifts by ≥0.2 to identify candidates. CIP: ciprofloxacin treated (50-65 µg/mL for the FQR strains or 0.11-0.12 µg/mL for WT). Untx: untreated. WT: wildtype. GPS: *gyrA parC adeS*. GPN: *gyrA parC adeN*. GPL: *gyrA parC adeL*.

**Figure 4:**
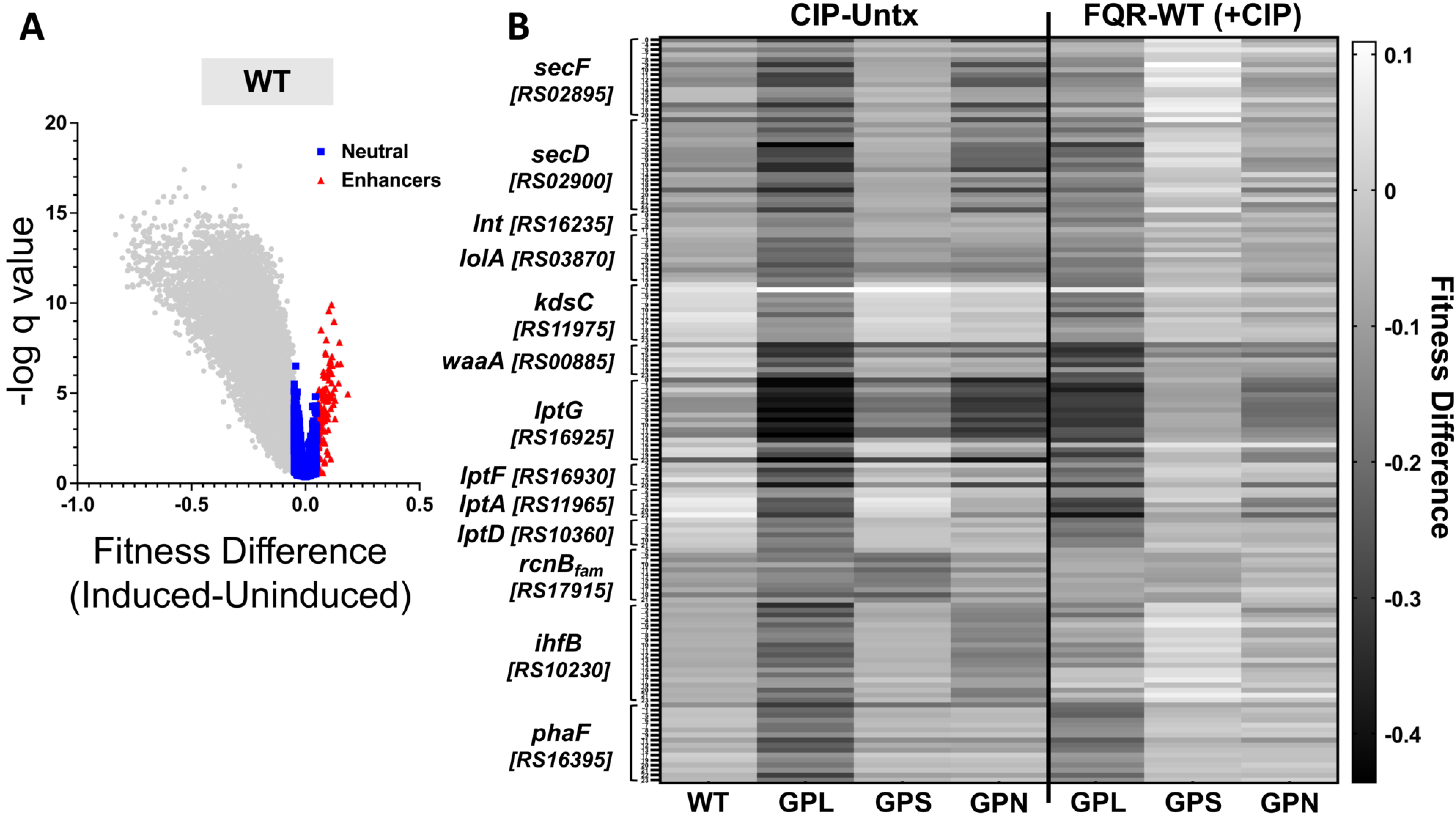
sgRNAs with minimal effects in the absence of drug that decrease fitness in the presence of CIP. (A) Statistical significance plotted as a function of mean fitness comparing growth in the presence and absence of dCas9 inducer (100ng/mL aTc) for 7-8 generations in the WT strain background (n=10). Gray: all sgRNAs. Blue: sgRNAs displaying neutral fitness effects (scores between -0.05 and 0.05). Red: sgRNAs revealing a fitness enhancement (scores of 0.05 or higher). (B) Heat map of neutral and enhancer sgRNAs (as defined in A) that significantly diminish FQR fitness in the presence of CIP. These pass the FDR of 5% (n=10) via multiple t-test analyses comparing hypomorphs incubated with and without drug (first 4 columns) and between strain backgrounds (FQR-WT) during CIP treatment (last 3 columns). Candidates were considered if at least 3 sgRNAs had fitness score decreased by ≥0.095 and at least one sgRNA did so in more than one FQR strain. Gene numbers are originally preceded by “ACX60_” for *A. baumannii* strain ATCC 17978UN (accession NZ_CP012004.1). 0: Perfect sgRNA. FQR: fluoroquinolone resistant. CIP: ciprofloxacin treated (50-65 µg/mL for the FQR strains or 0.11-0.12µg/mL for WT). Untx: untreated. WT: wildtype. GPL: *gyrA parC adeL*. GPN: *gyrA parC adeN*. GPS: *gyrA parC adeS*.

### Identification of hypomorphs with significantly altered fitness in the FQR strain backgrounds

To uncover factors that shape fitness of FQR pump hyper-expressing strains, multiple replicates (n=10) of libraries in four strain backgrounds (WT, GPS, GPN, and GPL) were cultured in rich LB broth in the presence of aTc inducer (for dCas9 expression), and fitness scores of hypomorphs were compared between the drug susceptible WT and FQR strains. To identify candidates important for maintaining FQR strain fitness under fluoroquinolone pressure, the libraries were also grown in parallel in the presence of sub-MIC levels of CIP at concentrations resulting in 30-40% increased doubling time across strains. Of note, there was minimal antagonism between aTc and CIP (Supp. Fig. 1D), and the effects of aTc on growth were similar for each of the strains. The statistical significance of each depletion (per unique sgRNA) was calculated as a function of the change in fitness between conditions or the difference in fitness between strains, determined by multiple t-test comparisons using an FDR or q value <5% (Materials and Methods) (33). Discoveries showing a significant fitness shift between comparisons were sorted *in silico* to identify sgRNAs yielding fitness scores at least 20% higher (knockdown imparting a fitness benefit or alleviating an effect) or at least 20% lower (knockdown causing a fitness defect or aggravating an effect). Among these, targets that were represented by 3 or more guides from their sgRNA set (with at least 10 reads at T1) were identified as top hits. Targets were identified that fulfilled these criteria during CIP treatment (vs untreated (untx)) and preferentially in the FQR strain (vs WT). Among these, the top candidates were targets identified in more than one FQR strain background.

In the absence of antibiotics, few candidates with three or more guides passed this statistical threshold, further supporting the Tn-seq results indicating that *A. baumannii* is quite tolerant of pump overproduction (Supp. Fig. 5A-C) (16). This was particularly true of the GPS strain, indicating that overproduction of the AdeAB pump is well tolerated by *A. baumannii* (Supp. Fig. 5A, vs. B and C). Loosening the fitness difference criterion (FQR vs WT) from at least 20% to at least 10% identified a total of 28 targets (originally 1) that reduced GPS fitness relative to WT, including proteins involved in secretion across the inner membrane (*secY, secA, ffh)*, DNA replication, the electron transport chain, and the TCA cycle (Supp. Fig. 5D-E, Supp. Fig. 6A). Doing so also uncovered 9 targets that diminished fitness of all three FQR strains (Supp. Fig. 5D) and 29 in at least two FQR strains (Supp. Fig. 5E).

**Figure 5:**
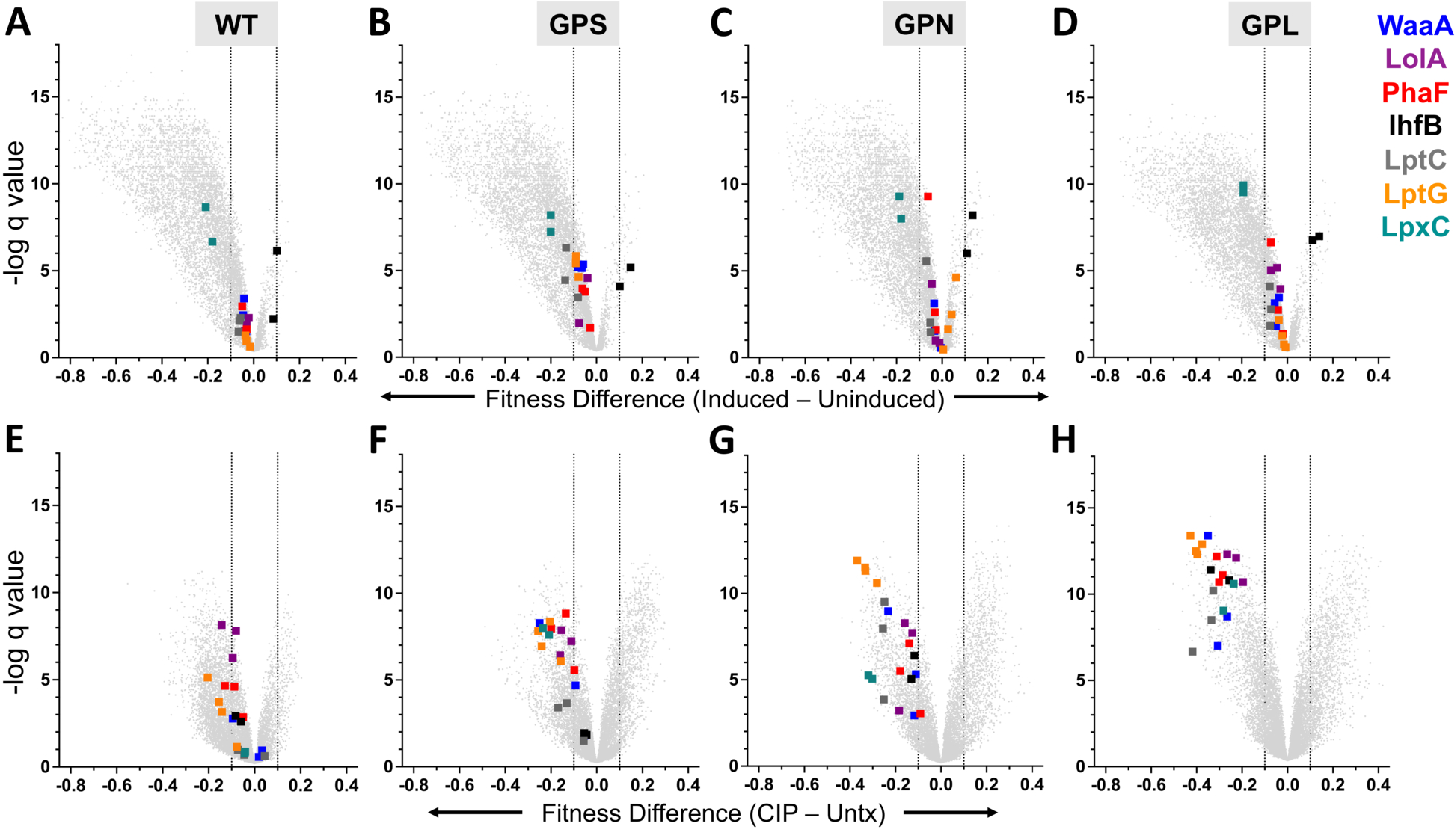
FQR strain hypomorphs that show high fitness levels in the absence of drug and are hypersensitive to CIP affect proteins critical to membrane integrity. (A-H) Statistical significance is plotted as a function of average fitness score differences for hypomorphs comparing dCas9 induced and uninduced conditions (A-D) or comparing CIP and Untx conditions (E-H) in the WT (A, E), GPS (B, F), GPN (C, G), or GPL (D, H) strain backgrounds (n=10). Gray: All sgRNAs. Seven prominent targets from the screen are highlighted, of which most depletions had little effect on fitness in the absence of drug but showed significantly lowered fitness in the FQR strains during CIP exposure. Dotted lines: selection parameters defining either fitness defect or high fitness. CIP: ciprofloxacin treated (50-65 µg/mL for the FQR strains or 0.11-0.12 µg/mL for WT). Untx: untreated. WT: wildtype. GPS: *gyrA parC adeS*. GPN: *gyrA parC adeN*. GPL: *gyrA parC adeL*.

**Figure 6:**
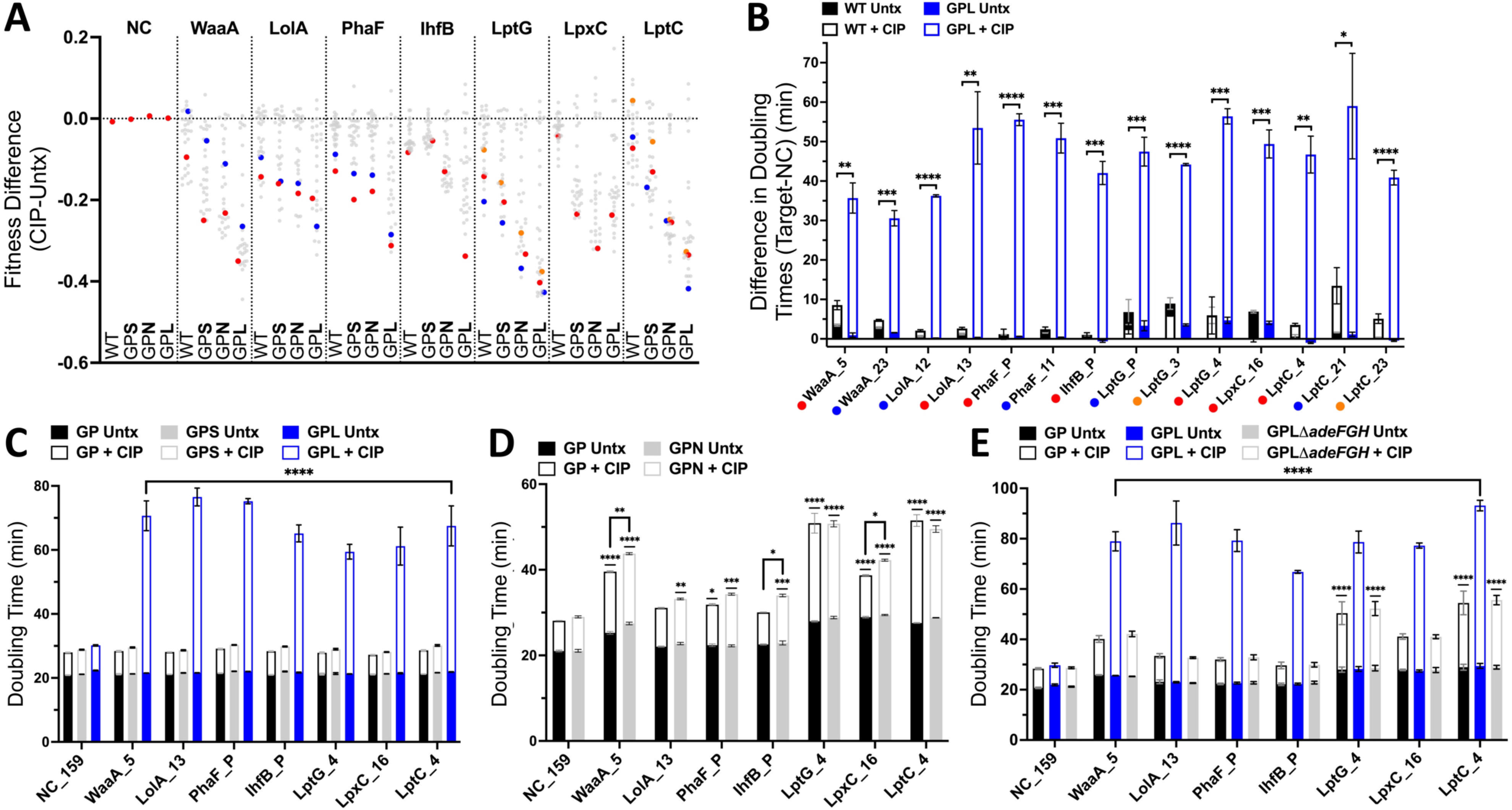
Individual CRISPRi validation assays confirm increased dependence of GPL and GPN strains on LOS biogenesis proteins for growth during CIP stress. (A) Mean fitness score changes imparted by each sgRNA during CIP exposure as compared to untreated conditions as derived from the CIP CRISPRi pooled screen (n=10). Dotted line represents the mean of scrambled controls that should exert no change in fitness. Light gray: all sgRNAs designed for the listed target (mismatched and shuffled guides included). Data points in red, blue, or orange highlight individual sgRNAs used to validate their respective targets’ impact on FQR fitness, as labeled in B. (B-E) These individual sgRNAs were constructed and hypomorphs in all strain backgrounds were cultured in the presence or absence of CIP. Doubling time differences in untreated and CIP treated conditions are superimposed. (B) Mean ± SEM (n=3) growth rates (in min) of hypomorphs are compared to those harboring a shuffle control sgRNA (NC_159). Unpaired t tests were applied with Welch correction and following the two-stage set-up of Benjamini, Krieger, and Yekutieli with a false discovery rate of 5% (33). Significant differences in doubling times yielded by targeted depletions between strains during CIP exposure are shown: *, p=0.063663; **, p<0.05; ***, p<0.01; ****, p<0.001. (C-E) Mean ± SEM (n≥3) growth rates are shown (in min) for the listed hypomorphs. Two-way Anova of pre-selected pairs with Šídák’s multiple comparisons revealed significant variations in growth rates between hypomorphs and the shuffle control within each strain background and between hypomorphs of two different strains (all compared to the GP parent), all during CIP exposure: *, p<0.05; **, p<0.01; ***, p<0.001; ****, p<0.0001. (C, E) Significant growth rate differences (****) were identified between all hypomorphs and NC in the GPL strain background as well as for all hypomorphs between the GPL and GP strain backgrounds. CIP: ciprofloxacin treated (using 50-65 µg/mL for the FQR strains or 0.11-0.12 µg/mL for WT in the pooled screen (A), while individual hypomorphs were verified using 0.1 µg/mL for WT (B), 10 µg/mL (E) or 13 µg/mL (C-D) for GP, 10 µg/mL for GPLΔ*adeFGH* (E), 50 µg/mL for GPS (C), 45 µg/mL for GPN (D), and 50 µg/mL (E) or 60 µg/mL (B-C) for GPL). Untx: untreated. WT: wildtype. GP: *gyrA parC*. GPL: *gyrA parC adeL*. GPN: *gyrA parC adeN*. GPS: *gyrA parC adeS*.

Outgrowth for additional generations revealed a smaller set of fitness determinants in GPS when compared to WT, only three of which overlap with those identified after 7-8 generations of growth (n=5, Supp. Fig 6B vs. Supp. Fig. 6A). However, it is notable that depletion of factors involved in competing processes resulted in opposing phenotypes after 14-16 generations. For example, depletion of the transcription termination factor Rho improved fitness of the GPS mutant relative to WT, while depletion of the anti-terminator NusG showed stronger fitness defects in GPS (*nusG* and *rho*; Supp. Fig. 6B). Importantly, when considering hypomorphs whose fitness varied by at least 20%, depletion of anti-transcription termination factor *nusG* was the single target identified as lowering fitness of more than one pump hyper-expresser in the absence of drug pressure. In fact, knockdown of *nusG* showed profound reductions in fitness in all three FQR strains when compared to WT (Supp. Fig. 5A-D). In *E. coli*, NusG-dependent antitermination is tightly linked to the regulation of ribosomal RNA transcription (34, 35).

### Depletions of OM integrity factors and F_1_F_0_ ATPase assembly proteins result in opposing effects

Using criteria similar to those used to identify strain-specific fitness differences in the absence of drug, we identified 8 targets with a fitness advantage and 10 candidates with a fitness defect in pump overproducers after exposure to CIP (Supp. Fig. 7). Analysis of pump overproducer and WT depletion derivatives (Figs 3A-F; Supp. Fig. 7A) revealed a clear pattern: the GPL background (Fig. 3C, 3F) generated the most depletions that selectively hypersensitized it to drug relative to WT, with few targets differentiating the GPS and WT backgrounds (compare Fig. 3D to 3E, F). This is consistent with a previous study analyzing mutations in nonessential genes (16). Furthermore, depletions leading to cell envelope integrity dysfunction were the most prominent candidates for CIP synergy, supporting the previous Tn-seq study (16).

**Figure 7:**
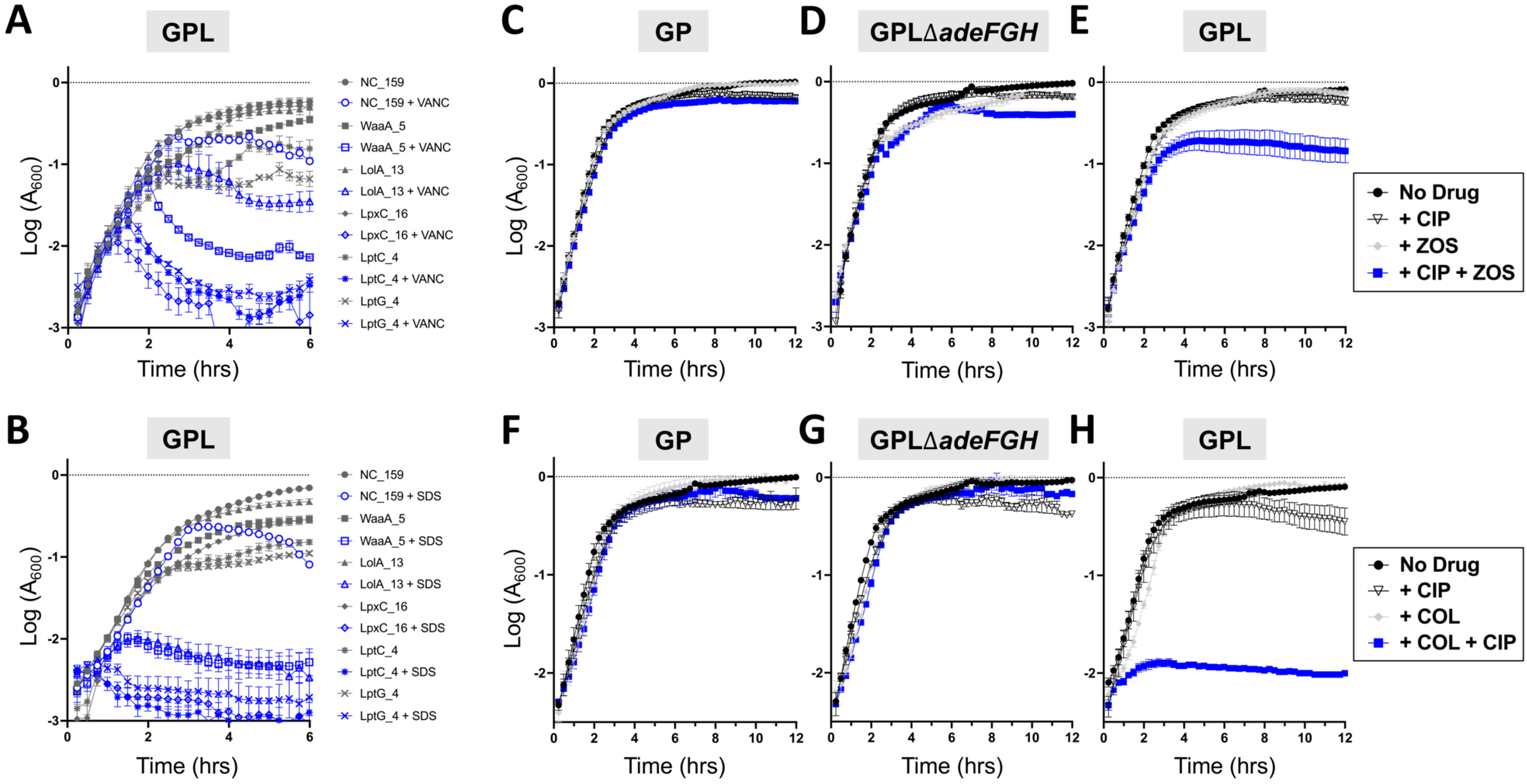
Depletion of LOS and lipoproteins results in defective envelope integrity, but ZOS and COL amplify CIP selectively in the GPL strain. (A-H) Growth of the indicated strains in LB at 37°C is displayed as the mean ± SEM (n≥3) after Y values were log transformed. (A, B) The indicated hypomorphs were grown in the absence of CIP with or without 64 µg/mL VANC (A) or 0.05% SDS (B), all in the GPL strain background. (C-H) Strains were incubated in the presence or absence of 2.5µM ZOS (C-E) or COL at 0.2 µg/mL (F-H) together with CIP at 13 µg/mL (C, D), 15 µg/mL (F, G), or 60 µg/mL (E, H). CIP: Ciprofloxacin. ZOS: Zosurabalpin. COL: Colistin. VANC: Vancomycin. SDS: Sodium dodecyl sulfate. GP: *gyrA parC*. GPL: *gyrA parC adeL*.

Depletion of *lpxC* and *lpxD* genes, encoding LOS biogenesis proteins, uniformly caused fitness defects relative to WT in the presence of drug, even for the robust *adeS* mutant (Fig. 3, Supp. Fig. 7B). With GPN and GPL, the spectrum of depletions that caused fitness defects was expanded to include other loci involved in LOS biosynthesis (*kdtA/waaA*, *kdsD/kpsF*) as well as a large cluster of genes involved in LOS transport to the surface (*lptA-C*, *lptG*) (Fig. 3, Supp. Fig. 7B). These data emphasize that OM integrity maintenance is required for FQR survival in the presence of drug, with factors controlling the composition of LOS being particularly important. Of note is the fact that depletions of BamD had weaker effects on the GPS pump overproducer than the other two FQR strains, consistent with our observation that there were fewer hypomorphs interfering with composition of the OM that reduced GPS fitness (Fig. 3, Supp. Fig. 7B). Notably, the strain analyzed in our studies is missing AdeC, the OM component of the *adeS*-regulated AdeAB system that is present in many other *A. baumannii* strains. Instead, AdeAB uses AdeK as the outer membrane component, which is not upregulated in response to *adeS-*activating mutations. We hypothesized that lack of OM component expression may be linked to resilience in the face of OM biogenesis defects, consistent with the general insensitivity of *adeS* mutants to outer membrane integrity defects (18, 36).

Our most surprising result was that depletion of loci associated with ribosome biogenesis and assembly, as well as translation, strongly increased fitness of at least two FQR strains relative to WT, consistent with slower translational rates resulting in increased fitness of efflux pump overproducers (Fig. 3 vs. Supp. Fig 7A, Supp. Fig. 7C). Of note, there were 13 (GPS) or 22 (GPN and GPL) ribosomal proteins that demonstrated higher fitness in the FQR strain backgrounds when depleted during CIP treatment compared to untreated, whereas 7 showed reduced fitness in the WT strain. Under CIP exposure, hypomorphs of 27 (GPS) or 46 (GPN and GPL) ribosomal proteins exhibited better fitness in the FQR strain backgrounds than in WT. In addition, depletions of *spoT* and *obgE*, which are predicted to cause translational stalling (37, 38), similarly increased the fitness of the GPL pump overproducer relative to WT (see Supp. Data 4). There was also increased fitness exhibited by depletions of the F_1_F_0_ proton-pumping ATPase/synthase in the presence of CIP, tying fitness to overall proton flux (Fig. 3, Supp. Fig. 7C). As the antibiotic efflux pumps act to antiport drugs in exchange for protons, decreased F_1_F_0_ ATP synthase activity may remove an important competitor for protons or alter cytoplasmic pH homeostasis in a fashion that favors drug efflux.

### High-fitness hypomorphs that are hypersensitive to CIP are largely impaired in envelope integrity

We next focused on guides that showed minimal fitness defects in the absence of antibiotic treatment, allowing condition-specific effects to be detected across a large dynamic range (Fig. 4A). This set of sgRNAs, called neutral or enhancer guides, were categorized as those resulting in fitness scores that averaged around 1.00 (neutral) or 1.06 (enhancer) during dCas9 induction in the WT background. They demonstrated no substantial change in fitness between growth in the presence or absence of inducer (fitness score differences within 0.05) (Fig. 4A). Of 284 targets that comprise the neutral and enhancer guide list, excluding shuffles and requiring representation by at least 3 unique sgRNAs, 13 candidates were identified that showed significantly lower fitness in the FQR background compared to WT in the presence of CIP (Fig. 4B, Supp. Data 4). These 13 candidates are predominantly proteins that shape OM integrity, with WaaA and Lpt proteins being top discoveries when restricting guides to those with inefficient knockdown (Fig. 4B).

Of the 13 FQR mutants that were sensitized to CIP, WaaA, LolA, PhaF, IhfB, LptC, LptG, and LpxC were selected for experimental verification. The sgRNAs were chosen based on their strong fitness defects during CIP exposure in GPL and at least one other FQR strain (Fig. 5B-D vs Fig. 5F-H), as well as having minimal effects on WT fitness in the absence or presence of drug (Fig. 5A, Fig. 5E). We observed the strongest fitness deficiencies in the GPL strain background (Fig. 5H), so we created individual sgRNA knockdowns of the top discoveries in GPL and evaluated their growth alongside those in the WT strain background.

Multiple sgRNAs were individually generated and tested for each target (highlighted in red, blue, or orange symbols, with their respective fitness defects during CIP exposure in the pooled screen shown in Fig. 6A). These hypomorphs were cultured separately in the presence and absence of CIP and their growth rates compared to a strain harboring a shuffled non-targeting sgRNA (“NC”, NC_159), growing for the same time periods as performed in the screen (Materials and Methods). The doubling times of hypomorphs were then compared to those of the control to detect altered growth rates caused by depletions (Fig. 6B, with the sgRNAs marked the same way as in Fig. 6A for reference). Depletion of WaaA, LolA, PhaF, LptG, IhfB, LptC, and LpxC all significantly slowed growth of GPL in comparison to WT during CIP exposure (Fig. 6B).

Notably, not all tested sgRNAs yielded the expected phenotype in the GPL background (Supp. Fig. 8A, marked by arrows), but this was linked to poor knockdown, as indicated by qRT-PCR (Supp. Fig. 8B) or poor growth in the absence of the drug. Given the latter reason, additional replicates of LptC and LpxC hypomorphs were tested. Depletions of LpxC and LptC resulted in heightened CIP sensitivity in GPL, with statistical significance only observed for LpxC (Supp. Fig. 9A). Finally, the top 13 candidate hypomorphic guide RNAs were introduced into the GP, GPS, GPN, and GPL strain backgrounds and then cultured individually in the presence or absence of CIP to determine allele specificity. Again, the GPL strain background showed an appreciably heightened sensitivity to CIP upon depletion of all targets of interest, but none significantly altered the growth rate of GPS (Fig. 6C). On the other hand, GPN, which showed more fitness dependencies than GPS (Fig. 5G vs Fig. 5F) exhibited significantly stunted growth compared to GP upon depletion of *waaA*, *ihfB*, and *lpxC* (Fig. 6D). The fact that there were fewer sensitivity loci and much smaller defects than identified for GPL is consistent with our previous analysis of nonessential genes targeted by transposon mutagenesis (16).

**Figure 8:**
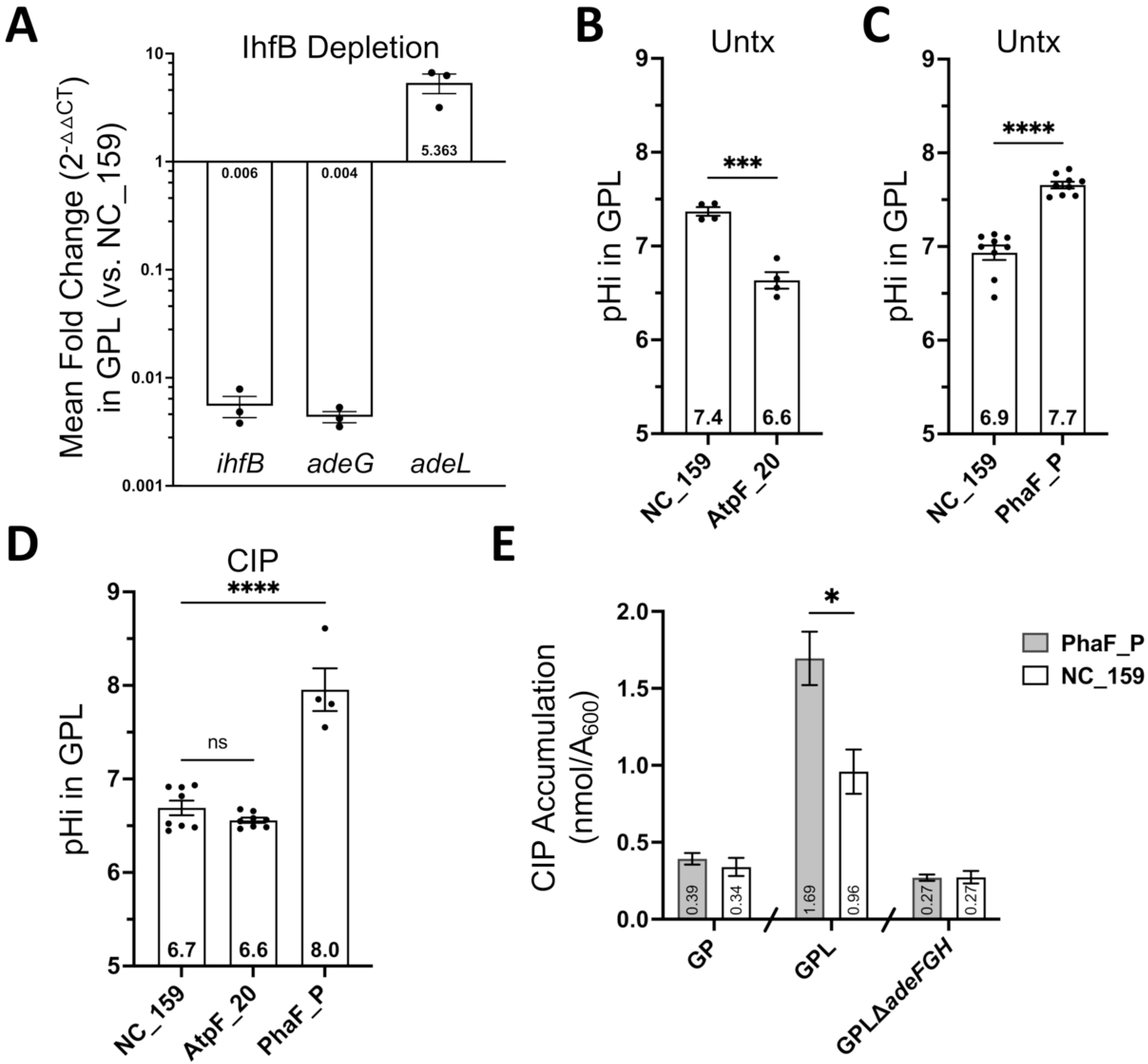
Non-OM pathways modulating fitness related to pump expression include the IhfB regulator and pH homeostasis. (A) Mean fold change (2^−ΔΔCT^) ± SEM (n=3) in transcript levels of each target in GPL during *ihfB* depletion with *ihfB*_P sgRNA relative to those observed with a non-targeting shuffle sgRNA (NC_159). (B, C) Mean ± SEM (n≥4) pHi values are displayed. Hypomorphs in the GPL strain background were incubated in LB with dCas9 inducer (100ng/mL aTc) to mid-exponential phase (A_600_∼0.455-0.638) before being loaded with BCECF-AM dye for cytosolic pH determination. Unpaired two-tailed t test: ***, p=0.0003; ****, p<0.0001. (B) One representative of two independent experiments. (D) Mean ± SEM (n≥4) pHi values are displayed for hypomorphs in the GPL strain background first incubated in LB with dCas9 inducer (100ng/mL aTc) to early mid-exponential phase (A_600_∼0.308-0.486) then for another 1 hour with CIP 60 µg/mL prior to BCECF-AM dye loading for pHi assessment. Ordinary one-way Anova with Dunnett’s multiple comparisons test: ns, adj. p=0.4729; ****, adj. p<0.0001. (D) One representative of two independent experiments for PhaF vs. NC after CIP exposure. (E) Mean ± SD (n=4) levels of intra-bacterial CIP accumulation shown for hypomorphs grown to early log phase then treated with CIP (13 µg/mL for GP and GPLΔ*adeFGH* or 60 µg/mL for GPL, concentrations that result in a 30-40% increased doubling time across strains) for 1hr. Unpaired t test with multiple comparisons between the shuffle control sgRNA (NC_159) and PhaF_P within each strain background corrected by two-stage step-up of Benjamini, Krieger, and Yekutieli with a FDR of 5%: *, p=0.000621. Untx: untreated. CIP: Ciprofloxacin. GPL: *gyrA parC adeL*.

To confirm that the defects linked to AdeL were the consequence of efflux pump overexpression, we challenged a derivative of GPL lacking AdeFGH (GPLΔ*adeFGH*) in the presence or absence of CIP (Fig. 6E). In the presence of an intact efflux pump, the doubling time escalated for hypomorphs in the GPL background, with growth rates significantly higher than those observed in the parent GP strain (Fig. 6E). Deletion of the AdeFGH efflux pump reversed this effect, consistent with the vulnerability associated with the *adeL* activation allele being attributed to pump overproduction (Fig. 6E).

### Depletion of LOS and lipoproteins impairs membrane integrity

It is anticipated that depletion of Lpt proteins, as well as other proteins involved in shaping the OM, disrupts membrane integrity. To test for OM dysfunction, hypomorphs were cultured in microtiter plates in the presence of vancomycin (VANC) or sodium dodecyl sulfate (SDS), and their growth was monitored in the absence of any other drug addition (Fig. 7A, B). As expected, depletion of WaaA, LpxC, LolA, LptC, and LptG yielded increased sensitivity to the addition of either vancomycin or sodium dodecyl sulfate, consistent with an impaired membrane barrier in these hypomorphs (Fig. 7A, B).

As depletions of LptC and LptG resulted in strain-specific fitness defects in the presence of CIP, we tested whether we could mimic genetic hypersensitization through the addition of zosurabalpin (ZOS), a drug targeting the LptB_2_FGC LPS transporter subcomplex (39). The strains GP, GPL, and GPL lacking AdeFGH were monitored for growth in LB with or without CIP and/or ZOS (Fig. 7C-E). ZOS in combination with CIP dampened the growth of the GPL strain, with the effect stronger than that observed for either drug alone (Fig. 7E). This aggravating effect was not observed in the GP parent strain (Fig. 7C), nor in the GPL strain lacking the AdeFGH efflux pump that is upregulated by the *adeL* mutation (Fig. 7D). An LOS-targeting drug colistin (COL) exhibited even stronger effects (Fig. 7F-H): COL in combination with CIP resulted in a drastic reduction of GPL growth (Fig. 7H). This effect was not detected in GP (Fig. 7F) nor in GPL lacking the AdeFGH pump (Fig. 7G). These results are consistent with the effects of Lpt, WaaA, and LpxC depletions (Fig. 6A, B) and illustrate that identification of mutations that cause aggravating effects also identifies drug targets that aggravate bacterial growth in the presence of a second drug.

### The IhfB regulator and pH homeostasis modulate fitness during CIP exposure to efflux pump overexpressers

The phenotype of *ihfB* hypomorphs was not directly linked to OM dysfunction, so we considered an alternative model for how they may impair fitness during CIP treatment. Given the documented role of IhfB in modulating transcription of numerous genes, including the AcrAB pump activator *robA* in *Salmonella* (40–43), it seemed likely that depletion of *ihfB* alters expression of key resistance determinants in *A. baumannii*. Therefore, transcript levels of *adeG*, encoding the RND protein of the AdeFGH efflux pump, and the *adeL* regulator were determined after IhfB depletion. While *adeG* was expressed about 920-fold higher and *adeL* about 5-fold higher in GPL than in GP (Supp. Fig. 10A), depletion of IhfB resulted in a steep decline in *adeG* transcript levels and a coinciding increase in *adeL* expression relative to the shuffle control sgRNA specifically in the GPL strain background (Fig. 8A, Supp. Fig. 10B). Therefore, reduction of pump gene expression resulting from IhfB depletion was positively correlated with decreased GPL fitness in the presence of CIP.

In addition to drug sensitizers, we identified hypomorphs that conferred a fitness advantage for efflux pump overexpressers in the presence of CIP. These included strains depleted of inner membrane-associated F_1_F_0_ ATPase/synthase subunits (*atpI, atpF* in all FQR strains; multiple other *atp* genes in GPL) (Fig. 3D-F). Some studies describe the bidirectionality of the F_1_F_0_ ATPase/synthase complex, having the ability to both synthesize and hydrolyze ATP using proton motive force (PMF) to drive ATP synthesis or reversibly hydrolyze ATP to pump protons out of the cytosol. ATPase activity, however, was found to be limited and structurally regulated in bacterial species such as *A. baumannii*, *E. coli*, and *M. smegmatis* (44–47). To determine if the F_1_F_0_ ATPase/synthase plays a role in maintaining pH homeostasis, intracellular pH (pHi) was measured during AtpF depletion in the GPL strain background. There was a significant decrease in cytosolic pH as a consequence of depletion (Fig. 8B), linking lowered ATPase activity to the failure to remove protons from the cytoplasm. This was true in the WT strain background as well (Supp. Fig. 10C). CIP treatment for 1 hour, however, eliminated the difference in cytosolic pH between AtpF-depleted and non-depleted cultures in the GPL strain background, indicating that alteration in pH homeostasis may not be a determinant that facilitates drug survival of the GPL strain after F_1_F_0_ ATP synthase depletion (Fig. 8D, Supp. Fig. 10D). Interestingly, AtpF depletion still significantly reduced the cytosolic pH of the WT strain (Supp. Fig. 10D).

Depletion of PhaF, which is a CIP sensitizer for the GPL strain, showed the opposite effect on cytosolic pH. This protein is part of the monovalent cation/proton antiporter PhaACDEFG complex, and it may modulate the proton gradient across the cytoplasmic membrane (48–51). When depleted of PhaF, both the GPL and WT strains showed increased cytosolic pH (Fig. 8C, Supp. Fig. 10C). An increase in both strains relative to shuffle controls was also observed after CIP exposure (Fig. 8D), albeit to a lesser degree in WT than in the GPL strain background (Supp. Fig. 10D). Although the exact pHi for the control varied across experiments, the effects of each hypomorph relative to control were reproducible across two independent experiments (Fig. 8B-D).

As depletion of the putative cation-proton antiporter increased sensitivity to CIP and disrupted cytosolic pH buffering, we tested the model that the PhaF protein is required for proper AdeFGH pump function. To measure AdeFGH pump activity, the accumulation of CIP was quantified in the GPL strain, the parental strain, and a GPL strain lacking the AdeFGH pump (Fig. 8E). To this end, each bacterial strain harboring either a PhaF knockdown plasmid or a nontargeting control was incubated in the presence of CIP at concentrations sufficient to increase doubling times by 30-40%. Intracellular CIP concentrations were then determined from extracts by mass spectrometry/liquid chromatography as described (77). The PhaF depleted GPL strain showed ∼50% increase in CIP accumulation compared to the shuffle sgRNA control (Fig. 8E). In contrast, PhaF depletion resulted in no alterations in CIP accumulation in either the GP strain or the GPL strain lacking the AdeFGH pump (Fig. 8E). Therefore, the absence of PhaF specifically interfered with AdeFGH pump function but otherwise had no effect on CIP accumulation.

## Discussion

Hyperexpression of RND drug efflux pumps often drives MDR, making them excellent targets for antimicrobial therapy. As essential proteins are particularly attractive targets for the development of new antimicrobials, we knocked down essential genes to identify hypomorphs sensitized to CIP in strains that specifically express fluoroquinolone efflux pumps (AdeFGH, AdeAB, and AdeIJK). This approach showed that depletion of transcripts encoding proteins involved in LOS synthesis and transport as well as lipoprotein maturation and trafficking to the OM all reduced the fitness of pump overproducers in the presence of antibiotic (CIP) treatment. Furthermore, CIP sensitivity was particularly pronounced in the GPL (AdeFGH pump-hyperexpressing) strain and was dependent on the presence of the efflux pump.

The most prominent classes of hypomorphs that enhanced sensitivity to CIP in the pump overexpressing strains were predicted to disrupt outer membrane (OM) integrity. Such targets include BamD, an OM component of the Bam complex necessary for OM proteins (OMPs) assembly (52). Interestingly, we previously showed that transposon knockouts of SurA, a non-essential OMP assembly chaperone, also decrease the fitness of the GPL strain (the FQR strain that overexpresses *adeFGH*) when CIP is present (16, 52–54). Depletions of BamD may interfere with the proper assembly or insertion of the pump OM factor (OMF) or other OMPs that maintain the OM barrier (18). Alternatively, a partially assembled Bam complex may specifically disrupt the cell envelope, resulting in combined defects of periplasmic precursor accumulation and loss of OMP content, leading to OM dysfunction. Similarly, depletions of lipoprotein maturation and trafficking proteins (*lnt* and *lolA*) sensitized efflux pump overproducers to CIP. BamD is one of multiple Bam lipoprotein components, so depletions of *lnt* or *lolA* could directly phenocopy defects in the Bam complex.

Defective LOS biosynthesis is the second example of OM disruption that leads to drug hypersensitivity in pump overexpressors. It is noteworthy that depletions of both LOS synthesis (*waaA*, *kds*, and *lpx* loci) and LOS transport (*lpt* loci) factors sensitized pump hyperexpressers to CIP. Two earlier reports regarding drug hypersensitivity in *A. baumannii* showed fitness defects that resulted preferentially from defects in LOS transport (24, 55). Decreased levels of Lpt proteins produce a defective cell envelope that results in accumulation of material in the periplasmic space (56, 57). Presumably, the accumulation of LOS precursors inserted into the inner membrane in the absence of Lpt function results in a general sensitivity to most clinically effective antibiotics (24). In contrast to these results, when we focused specifically on strains expressing RND efflux pumps, the spectrum of sensitizing targets was broadened to LOS synthesis (*lpx*), and possibly any factor that disrupts OM biogenesis.

Defective envelope biogenesis does not explain why fitness is more severely compromised in the high-level FQR pump hyper-expressers than in the WT or GP strains. One explanation is that LOS directly binds to the RND transporter complex, and the absence of this lipid destabilizes the OM component of the transporter. It is known that LPS can bind OMPs and that OMP-interfacing lipids may mediate protein self-association, contributing to OM integrity (58, 59). By this model, proteins that shape the OM support assembly of the pump complex and thereby resilience in the presence of drug. Overall, the targets revealed by this work indicate that the FQR pump hyper-expressers are more vulnerable to changes in OM composition than strains lacking these pumps. Components that support the integrity of the permeability barrier and stabilize RND drug transporters are likely to be important for survival during drug treatment. Consistent with these results is the demonstration that RND pump overexpressing strains are hypersensitive to ZOS and COL, and compound P2-56-3 described in another study, all of which disrupt the OM (55).

In addition to targets linked to OM biogenesis, loci were identified that had no clear connection to supporting envelope integrity, but which also sensitize the pump overexpressing strains to CIP. These include IhfB and PhaF. IhfB is part of the nucleoid-associated integration host factor complex that can modulate chromosome structure and influence gene expression (40–43). Depletion of IhfB resulted in reduced expression of *adeG* accompanied by increased expression of the pump regulator gene *adeL*, consistent with low pump concentrations leading to CIP sensitization. Although it is unclear how a deficit in IhfB caused this defect, this result opens the door to targeting global regulators as a strategy for combatting drug resistance.

Perhaps the most profound defect found in this work was the demonstration that depletions of PhaF directly interfered with the ability of the AdeFGH pump to prevent cytosolic CIP accumulation. This protein is part of the monovalent cation/proton antiporter PhaACDEFG complex (48–51), which may control ion or pH homeostasis in the FQR strains to compensate for the increased drug-proton antiport activity of the RND efflux pumps. Upon depletion of PhaF, we observed increased cytoplasmic pH. In addition, LC/MS analysis of cytosolic extracts demonstrated that PhaF depletion resulted in increased accumulation of CIP when compared to the shuffle control sgRNA in the pump overexpressing GPL strain. Therefore, the Pha complex is required for proper activity of the AdeL-regulated pump, possibly playing a buffering role important for RND pump activity.

Depletion loci that resulted in enhanced fitness specifically in pump overexpressers were also identified after CIP treatment. These included the inner membrane-associated F_1_F_0_ ATPase/synthase subunits (*atpI, atpF* in all FQR strains and multiple other *atp* genes in GPL) and translation-associated factors (*fmt*, *rimM*, *der/engA*, *rnpA*). AtpF encodes the ATPase B subunit that connects the F_1_ catalytic domain to the F_0_ membrane-spanning proton translocator, linking proton translocation to ATP generation by F_1_ (46, 47). AtpI, on the other hand, functions as a poorly conserved non-structural chaperone, supporting the formation of the F_0_ c-ring and complex stability (60, 61). Depletion of the specific subset of components leading to drug tolerance may be a consequence of their limited lethality after depletion as compared to other components.

There are two clear models for how depletion of F_1_F_0_ ATP synthase components could provide a fitness advantage in the presence of the drug. First, lowering the cytosolic pH could simply reduce antibiotic efficacy (62, 63). Secondly, similar to the F_1_F_0_ ATP synthase, RND pumps utilize the proton motive force generated by the electron transport chain, and reducing ATPase synthase concentrations on the membrane may facilitate kinetic linkage of the efflux pump to the respiratory chain. This model is consistent with our previous observation that there were significantly reduced numbers of viable transposon insertion disruptions in the cytochrome b_o_ (3) ubiquinol oxidase complex in the GPL strain (16). Therefore, the oxidase complex could provide the protons necessary for efflux pump activity. In fact, depletion of the ubiquinol oxidase complex as well as other systems involving the electron transport chain reduced fitness of GPN and GPL even in the absence of drug in this study, indicating that the oxidase may play an important homeostatic role even in the absence of known substrates (Supp. Fig. 5E). Therefore, the ATP synthase and the ubiquinol oxidase appear to play opposing roles in supporting fitness of pump overproducing strains.

Interestingly, in our RNA-seq experiments, we found that *atpE* transcription was significantly downregulated in the GPL efflux overproducer when compared to the GP strain grown in the absence of drug (16). Although depletion of this critical F_0_ component was not shown to protect from drug action, it does emphasize the similarity between drug stress and stress related to overproduction of the AdeFGH pump. Manipulation of the proton motive force may be a critical determinant in protecting against both pump overproduction and drug exposure.

The ATP synthase depletion phenotype after CIP treatment parallels that observed for depletion of translation apparatus components. Translational disruption may be similar to ATP-depletion, as each assumes dormancy conditions that protect against drug stress. Both lowered ATP and reduced translation have been associated with persister cell formation (64, 65). Interestingly, pump hyperexpression has also been documented to contribute to the development of persistence (66), most notably in the case of the AdeFGH efflux pump (67). Therefore, either depressed ATP synthesis or translation could support the fitness of pump overexpressing isolates, allowing survival in the presence of high antibiotic levels. This may be one factor in explaining why translation inhibitors have occasionally been linked to antagonism of fluoroquinolone action (65, 68–71). In conclusion, this study exposes essential gene targets that enhance FQR strain sensitivity to CIP as well as targets that improve FQR strain fitness during drug exposure. Future efforts will be directed to exploring our hypotheses in hopes of better understanding both the hidden susceptibilities of pump over-expression and identifying proteins that help support the viability of the MDR state.

## Acknowledgements

This work is supported by NIAID grants U01AI124302, U19AI158076, R21AI128328, U19AI142780, and U19AI189168. We thank Megan W. Tse (Broad Institute of Massachusetts Institute of Technology and Harvard) for LptC and LptG hypomorphs, Defne Surujon (Boston College) for a sam-to-wig file conversion script, Yunfei Dai (Northeastern University) for communications regarding dCas9 strain generation, sgRNA design, and TRANSIT, and for the pYDE009 and pYDE007 plasmids, Rebecca Batorsky (Tufts Technology Services) for aiding with TRANSIT accessibility, Albert Tai (Tufts Genomics Core Facility) for communications regarding sequencing and library preparation, as well as Amir George (Rutgers University) for communications regarding their intra-bacterial drug accumulation assay.

R.R.I and E.H conceptualized and devised the study. E.H and W.H designed the sgRNA library. E.H, K.N, and I.N ran the preliminary screens with WT and GPS in the absence of drug. E.H executed the CIP screen and hit validations. W.H tested depletion efficiency of select high fitness perfect sgRNAs, provided bioinformatic support, and reviewed the manuscript. S. and J.S.F. performed and analyzed results of the intra-bacterial drug accumulation assay. E.G shared all strains, vectors, and methodologies for CRISPRi set up and edited the manuscript. E.H and R.R.I interpreted the results and wrote the manuscript.

## Materials and Methods

### Bacterial strains and growth conditions

Bacterial strains used in this work are listed in Supplementary Data 1. All *Acinetobacter baumannii* strains are derived from ATCC 17978UN. Cultures were grown in lysogeny broth (LB) (10 g/L tryptone, 5 g/L yeast extract, 10 g/L NaCl) or on LB agar plates (LB supplemented with 15g/L agar). Superoptimal broth with catabolite repression (SOC) was used for recovery following all electroporations of *A. baumannii.* Electroporations were performed using 0.1cm gap cuvettes and Bio Rad Gene Pulser, with the settings 200 Ω, 25 µF, and 1.8 kV. For growth in broth culture, bacteria were aerated at 37°C on a platform shaker, roller drum, or shaking in 96-well plates (Costar) with growth monitored by measuring absorbance via a spectrophotometer at 600nm (A_600_) with Genesys 30 (Thermo Scientific) or Epoch 2 or SynergyMx (Biotek) plate readers. Cultures were first incubated to early post-exponential phase in LB before diluted to a starting A_600_ of 0.003 in 5mL LB broth or in 200µl LB (96-well plates) supplemented with aTc for dCas9 induction (when hypomorphs are evaluated) and drug where applicable. Growth was monitored every 15 minutes (plate reader) or evaluated after 170min (dCas9 induced but not drug treated) and 230min (dCas9 induced and drug treated). LB was supplemented with antibiotics, as needed, at the following concentrations: aTc: 100ng/mL, Carbenicillin (Carb): 50/100µg/mL, Gentamicin (Gent): 10µg/mL, Chloramphenicol (Cm): 10/25µg/mL, VANC: 64 µg/mL, SDS: 0.05%, COL: 0.2µg/mL, CIP: varying concentrations (Sigma), or ZOS: 2.5µm (MedChemExpress).

### Molecular cloning and mutant construction

Primers (IDT or GENEWIZ) and plasmids used throughout this study are listed in Supplementary Data 1. The dCas9 construction is tetracycline (Tet)-inducible and chromosomally embedded at the Tn7 site in the four strains of interest. The plasmids and techniques used to create these strains are as previously described (26, 31). Briefly, the mobile Tet-inducible dCas9 was introduced into the four *A. baumannii* recipient strains of interest via four parental mating and Tn7 transposition. The recipient *A. baumannii* strains, the HB101 RK600Cm^R^ helper, EGE187 DH5α λpir (pTNS3) encoding the T7 transposase, and the YDE009 DH5α (pUC18T-miniTn7T-GentR with *tetR*-*P_Tet_::dCas9-rrnB*T1-T7Te) donor strain were grown overnight in LB with appropriate antibiotics. The following day the cultures were normalized to A_600_ of 2.0 and 100µl of each of the three *E. coli* cultures were combined with 100µl of each *A. baumannii* recipient culture in 600µl LB, prewarmed to 37°C. The mixture was washed twice in 37°C LB, then resuspended in 25µl of fresh broth and spotted on pre-warmed LB agar plates. These plates were incubated at 37°C for approximately 5 hours then resuspended in 1mL sterile phosphate buffered saline (PBS) before plating and overnight selection on Vogel-Bonner minimal medium (VBM) containing 10µg/mL of Gent to select against *E. coli* strains and the parental Gent^S^ *A. baumannii*. Exconjugants were colony-purified on VBM containing 10µg/mL Gent, and screened on LB agar with 100µg/mL Carb and on LB agar with 10µg/mL Gent. The insertion at the *att*Tn7 site downstream the *glmS* gene was confirmed by colony PCR using the primers glmS2-F1 and Tn7R-R1.

The vector pYDE007 (Addgene plasmid # 194152; http://n2t.net/addgene:194152; RRID:Addgene_194152) used to introduce the sgRNAs is a shuttle plasmid previously described (26) containing the origin of replication of pBR322 and an origin of replication from a natural *Acinetobacter* isolate pWH1277 (26) to allow for propagation in both *E. coli* and *A. baumannii* respectively. It encodes for the β-lactamase gene (*bla*) on the backbone and contains a sequence to be replaced by an sgRNA sequence upstream of the dCas9 handle and *S. pyogenes* transcriptional terminator, with the fusion driven by the constitutive synthetic promoter pJ23119. For efficient cloning of sgRNAs, the vector was made GoldenGate^TM^-compatible by first mutating the single BsaI Type II restriction site present in the *bla* gene and then introducing two BsaI sites between the promoter and dCas9 handle to allow for directional sgRNA oligo insertion (pEHE51, Addgene plasmid # 250522). Using this backbone, all 24nt sgRNA oligos were constructed with additional 4nt overhangs that allowed for directional cloning. The sgRNA bank was designed by loosely following a previous strategy for stoichiometric knockdowns (20), and the vector and library design were used for construction of the sgRNA-plasmid library by contract (GENEWIZ).

### Identification of essential genes in *Acinetobacter baumannii*

Essential genes of *A. baumannii* ATCC 17978UN were determined using transposon mutagenesis sequence data in the WT strain (16, 25), and the data from that study was subjected to TRANSIT Gumbel analysis to identify the essential genes (26–28). Briefly, sequence mapped SAM files were converted to wig files with a custom python script bowtie_to_wig2.py provided by Defne Surujon from the Tim van Opijnen lab. The wig files were then used as input, using the TRANSIT Gumbel analysis pipeline (28) with default settings. Cross referencing our list of essential genes to those similarly identified in other studies in the same strain or the more recently isolated AB5075 clinical strain revealed 406 genes exhibiting a high likelihood of essentiality, given their classification as essential in at least one of the other published analyses as well (26, 29, 30).

### sgRNA library design

The 24nt sgRNAs were principally designed (to target each essential gene) with the antisense strand sequence using a PAM site on the antisense strand that is as close as possible to the translational start site. The sgRNA pool was then expanded with an additional 23 mismatch sgRNAs that differ from each targeting sgRNA by 1 or 2 nts, along with 5 shuffle sgRNAs that comprise the same nts as the original target-homologous sgRNA but are shuffled to minimize the likelihood of off-target effects. An additional 216 “pool shuffle” sgRNAs of an array of nts with the GC content of the targeted genes designed to minimize homology to any genomic sequence, bringing the final sgRNA pool to 12,000 sgRNAs total (Supp. Data 2). Mismatches were employed to introduce sgRNAs with reduced knockdown efficiencies, while the shuffles served as negative controls (anticipated to provide a neutral effect on fitness). A BLAST search did reveal that up to 12bp overlap with the genome was unavoidable among some members of the control pool. These mismatches and shuffle sgRNAs were designed as previously described (20), using the python script ‘final sgRNA dirty pool design.ipynb’ (accessible via GitHub: https://github.com/isberglab/CRISPRiaDesign.git).

### Preparation of pools with sgRNA library

The sgRNA-pDNA library was introduced into the FQR pump hyper-expressers and WT strain with chromosomally encoded Tet-inducible dCas9 via electroporation. About 50µl electrocompetent cells were transformed with 23-46ng of the pDNA, the resuspended mixture diluted 1/10, and 200µl plated on five large pre-warmed SOC agar plates overlain with mixed cellulose ester filters (0.45µm pore size, 137mm diameter, Advantec). The mixture was spread on the filters using beads. This was repeated for a total of 3 transformations per strain background. These transformants were allowed to recover at 37°C for an hour on SOC agar plates before the filters were transferred to selective LB agar plates containing 50µg/mL Carb and incubated overnight at 37°C for about 13.5-14 hours. The following day, filters were pooled to acquire somewhere around 250,000-350,000 CFUs, estimated based on using image analysis of previous pools (ImageJ). Pooling entailed washing colonies off with 100mL sterile PBS with gentle shaking in a sterile round bottom 3L beaker. Glycerol was added to a final concentration of 10-20% and 1mL aliquots were placed into cryotubes for -80°C storage until needed.

### Identification of hypomorphs with fitness defects

Following previous protocols for determining fitness using Tn-Seq pools (10, 25), each vial of *A. baumannii P_Tet_::dCas9* harboring the sgRNA array was thawed on ice, diluted into 25mL LB to an A_600_=0.1, and revived by incubation at 37°C with shaking for 50min-60min until they reached around A_600_=0.2. Each culture was then diluted to an A_600_=0.003 in 10mL LB in the absence or presence of 100ng/mL aTc for dCas9 induction and the fluoroquinolone CIP. 100ng/mL of aTc inducer was used to achieve comparable levels of dCas9 expression in each strain background (approximately 20-30X induction) while minimizing growth inhibition (equivalent to a 9.5-11.2% increase in growth rate compared to uninduced in all four strain backgrounds in the absence of CIP). Cultures were then treated with CIP concentrations yielding a 30-40% increase in growth rate relative to those grown in parallel with inducer in the absence of CIP. Cultures were incubated for approximately 7-8 generations on the drum wheel at 37°C, for about 170 (no drug treatment) or 230 (CIP treated) minutes, until densities reached A_600_= 0.5-0.895. The cultures were harvested both before (T1) and after (T2) dCas9 induction and drug treatment to allow sgRNA fitness analyses. Sample pellets were stored at -20°C until all were collected. Plasmid was extracted from each sample using the QIAprep Spin Miniprep Kit (Qiagen) following manufacturer’s protocol.

sgRNAs were amplified from harvested plasmid DNA using primers For2_sgRNA_Pool and Rev_sgRNA_Pool at 0.6µM in a 50µl PCR reaction with 10µl 5x Q5 buffer, 0.5µl of Q5 hot start polymerase, 1µl 100mM dNTPs (NEB), 100ng of the pDNA, and UltraPure water (Thermo Fisher Scientific). The thermocycler settings were as follows with a heated lid: 98°C for 1min, 14 cycles of 98°C for 10sec, 70°C for 20sec, and 72°C for 1min, then finally 72°C for 2min. Barcodes were then introduced with a second round PCR using a legacy Nextera kit of adaptors (Illumina, compatible with FC-121-1031, FC-121-1030, FC-121-1012, FC-121-1011). The second PCR was performed using 1µl of the first PCR product, 4µl UltraPure water, 7.5µl 2x NEBNext High Fidelity Master Mix (NEB), and 1.25µl of each barcoded Nextera N5x and N7x index primers per reaction. The thermocycler was set up to hold a heated lid while running at 72°C for 3min, 98°C for 2:45min, and 8 cycles of 98°C for 15sec, 65°C for 30sec, and 72°C for 1:30min. The desired amplified product is 328nt in length. 5µl samples of the PCR2 products were fractionated on a 2% agarose Tris-Acetate-EDTA (TAE) gel using Sybr Safe dye (Invitrogen) alongside a set amount of 100bp DNA ladder (NEB) and run for 1hr at about 90V for quantification. Samples were imaged and concentrations measured using a BioRad quantitative gel doc system, with quantitation performed using Image J, or dedicated Bio-Rad software so that approximately 100ng of each sample was pooled for sequencing. The samples were PCR column purified (Qiagen) and their final concentrations determined via Nanodrop prior to submission to Tufts University Core Facility of Genomics for sequencing. The pools were sequenced via Illumina HiSeq 2500 SE100 using a legacy Nextera primer mixture.

The 5’ sequence preceding the 24nt sgRNA was trimmed using Cutadapt (72). Reads were assigned to each sgRNA using a Python script as described (21). To better assess fitness of all hypomorphs in the pool, a set of 25 shuffle sgRNAs were chosen to serve as negative/neutral controls: The dataset used was from a preliminary study comprising 10 replicate cultures from the WT and GPS strains harboring the sgRNA library grown in rich media in the presence or absence of dCas9 induction. To identify the neutral controls, the assigned reads for all sgRNAs at T1 and at T2, in the induced and uninduced cultures in each strain background, were normalized to the total reads in each replicate sample and then transformed by 1x10^6. As there were 10 replicates per condition, these values were averaged across replicates for each sgRNA. The median was then determined for each condition and the sgRNAs falling within the 25-75th percentile were identified. Shuffle sgRNAs that fell in this range across all conditions in both the WT and GPS strain backgrounds were identified as candidates for the normalization controls. Among these sgRNAs were a set of guides that also yielded mean fitness values within the 25th-75th percentile in more than one condition, 25 of which were selected as neutral controls for downstream fitness normalization.

To calculate fitness of all hypomorphs screened in the pool, read counts for each sgRNA were first normalized to the total assigned reads per sample. Any counts of 0 were given a read count of 1 for downstream fitness calculations (32, 73). Fitness values were then normalized to the set of 25 shuffle sgRNAs serving as neutral controls by dividing each individual sgRNA fitness score by the average fitness of those 25 sgRNAs in each sample. Normalized fitness values of each sgRNA were then compared as described in Results. Multiple unpaired t-test analysis was used to identify significant differences in fitness as a consequence of drug treatment or strain background. Discoveries of fitness scores that significantly differ in these comparisons were determined by the two-stage step-up method of Benjamini, Krieger, and Yekutieli with Q < 5% (33). Findings characterized as top discoveries were targets represented by 3 or more sgRNAs. Those with <10 cumulative reads across replicate cultures at T1 prior to outgrowth as well as ribosomal proteins were excluded. Data pertaining to the CIP screen are shown in Supplementary Data 4. All sequence data is also deposited in the SRA database under BioProject number PRJNA1067098.

### Validation of individual CRISPRi fitness defects

Individual sgRNA vectors were constructed by digestion of the pEHE51 vector with BsaI-HFv2 (NEB) prior to gel extraction followed by guide oligonucleotide insertion. The 24nt oligonucleotide guides (1µl of each complementary oligo at 100µM) containing 4nt overhangs were incubated with T4 PNK (NEB) for 1.5hrs at 37°C, followed by addition of 2µl 1M NaCl and oligo annealing starting at 95°C, with ramping to 25°C performed using a temperature drop of 15-20°C every 5min. Oligos were diluted 100X and 4µl were ligated to 100ng vector with T4 DNA Ligase (NEB) overnight at 16°C prior to electroporation into dCas9-harboring strains. Individual hypomorphs were tested by growth to early post-exponential phase in LB, followed by dilution to A_600_∼0.003 in LB supplemented with 100ng/mL of aTc with or without CIP (at varying concentrations) and measuring A_600_ at 170 min (untreated) or 230 min (CIP treated). Doubling times were determined (Time_min_ x Log(2))/(Log(A_600@T2_)-Log(A_600@T1_)).

### pHi Assay

This assay was performed as previously described (16), adapted from earlier work (74–76). The hypomorphs (and shuffle controls) evaluated in this work were grown to late exponential phase before dilution to A_600_ = 0.01 in 8mL LB + 100ng/mL aTc. These cultures were incubated to mid-exponential growth phase without additional drug (Untx) or to early-exponential growth phase then treated with CIP (0.1 µg/mL for WT or 60 µg/mL for GPL) for an additional hour. About 2.5-5mL of bacteria were harvested (approximately A_600_ ∼1.5-3.5 units) then re-suspended in 1mL PBS. Per sample, 50-100µl of culture in PBS was incubated with 50-100µl of PBS pre-mixed with 40µM BCECF-AM dye (1:1) for a final concentration of 20µM BCECF-AM dye (Thermo Fisher Scientific) in a 96-well v-bottom plate. They were incubated for 30min at 30°C, pelleted, energized in 200µl LB for 5min at 37°C, then pelleted and re-suspended in 210µl mixed dibasic sodium phosphate and monosodium phosphate buffer of varying pH (standard curve) or neutral pH (experimental samples). 100µl of each sample was aliquoted by multi-channel into a black with a clear flat bottom 96-well plate pre-loaded with 100µl buffer (and 10µM CCCP in the wells designated for the standard curve) for downstream pHi evaluation via 490/535 and 440/535 excitation/emission detection.

### Intra-Bacterial Drug Accumulation Assay

Hypomorphs (and shuffle controls) were cultured to early-exponential phase from a starting A_600_ = 0.01 in LB broth with 100ng/mL aTc for dCas9 induction. Cells were then treated with CIP (13 µg/mL for GP and GPLΔ*adeFGH* or 60 µg/mL for GPL) for 1 hour, then 0.5 A_600_ units were pelleted and frozen. These samples were processed for LC-MS intra-bacterial quantification of CIP as previously described (77).

### qRT-PCR

Cultures were grown in LB broth to early post-exponential phase before dilution into 5mL LB with 100ng/mL aTc to a starting A_600_ of 0.003 (A_600_ of 0.01 in 3mL total volume for AtpF depletion). Cultures were harvested after 170min reaching A_600_ of 0.134 to 0.904 (average 0.466). The following steps are as previously described (16): RNA was extracted using RNeasy (Qiagen). RNA was treated with ezDNase before cDNA synthesis with the SuperScript VILO cDNA kit (Thermo Fisher). Targets were amplified using PowerUp SYBR Green Master Mix (Applied Biosystems) and CT values determined using the StepOnePlus Real-Time PCR system (Applied Biosystems). The efficiency of primer pairs used for target amplification was evaluated using a standard curve of cDNA at varying dilutions and consequently verified to be about 94.3-113.5% for each pair used and within 10% efficiency of 16S control. Three biological replicates were evaluated for each strain or hypomorph and three technical replicates examined per biological sample during target amplification. To test for gDNA contamination, a set of control samples were tested in parallel which were not reverse transcribed during the cDNA synthesis reactions. Transcript levels of each specified target were compared between each hypomorph and the control strain harboring a non-targeting shuffle sgRNA, NC_159, after normalization to levels of 16S.

